# Assessment of microbial communities in Nakivubo wetland and its catchment areas in Lake Victoria Basin, Uganda

**DOI:** 10.1101/2020.12.08.413922

**Authors:** Bernard N. Kanoi, Maribet Gamboa, Doris Ngonzi, Thomas G. Egwang

## Abstract

Microbial community structure changes are key in detecting and characterizing the impacts of anthropogenic activities on aquatic ecosystems. Here, we evaluated the effect of river pollution of industrial and urban sites on the microbial community composition and distribution in the Nakivubo wetland and its catchment areas in Lake Victoria basin, Uganda. Samples were collected from two industrial and one urban polluted sites and the microbial diversity was analyzed based on a 16S rRNA gene clone library. Differences in microbial diversity and community structure were observed at different sampling points. Bacteria associated with bioremediation were found in sites receiving industrial waste, while a low proportion of important human-pathogenic bacteria were seen in urban polluted sites. While *Escherichia* spp. was the most dominant genus of bacteria for all sites, three unique bacteria, *Bacillus sp., Pseudomonas sp., Thermomonas sp*., which have been reported to transform contaminants such as heavy metals and hydrocarbons (such as oils) by their metabolic pathways were also identified. Our results may serve as a basis for further studies assessing microbial community structure changes among polluted sites at Nakivubo Water Channel for management and monitoring. The diversity of natural microbial consortia could also be a rich bioprospecting resource for novel industrial enzymes, medicinal and bioactive compounds.

## Introduction

Industrial and residential development has introduced a wide range of environmental pollutants into freshwater ecosystems on a continual basis, impacting the biodiversity on a global scale (e.g. Dudgeon et al. 2006), and affecting human health (e.g. Haseena et al. 2017). Anthropogenic pollution can cause disproportional dispersal of pollutants into freshwater ecosystems strongly influencing prolonged changes on the aquatic biota.

Microorganisms represent the most useful indicator of the biological quality of freshwater ecosystems (Paerl et al. 2003). Their abundance and functional diversity is strongly affected by water pollution (Chakraborty and Bhadury 2015; Cabral 2010). Microorganisms have become an important component of examining pollution of freshwater ecosystems by anthropogenic activities mainly due to their direct responses to environmental alterations (Upadhyaya and Verma 2015). Specifically, microbial communities have shown a wide range of metabolic diversity capacity by using different organic and inorganic pollutants that allow them to tolerate and adapt to polluted environments thereby making them effective agents for bioremediation (e.g. Coelho et al. 2015; Chakraborty and Bhadury 2015). However, the complex effect of the pollutants on freshwater ecosystems can lead to changes in microbial community composition and diversity and an alteration of community functionality (Han 2006; Colin et al. 2012; Gilbert et al. 2012).

The advance of novel tools to monitor and characterize microbial community diversity at the molecular level has facilitated the tracking of microbial community responses to pollutant exposure with high efficiency (Franco-Duarte et al. 2019). Among the molecular approaches, 16S ribosomal RNA gene clone library has been one of the most frequently used techniques for taxonomic identification and characterization of microbial organisms (e.g. Huang et al. 2013; Warnecke et al. 2004). Here, we investigated the effects of industrial and urban pollution on the diversity and distribution of freshwater microbial communities at Nakivubo wetland and its catchment areas in the Lake Victoria basin of Uganda.

Nakivubo Water Channel has a 50 km^2^ catchment area, covers 12.3 km and feeds the wetlands of Kampala city, Uganda (Emerto 1998). The channel receives a lot of the domestic and industrial wastewater of the central division of Kampala (Matagi 2002). Previous studies showed high concentrations of thermotolerant coliforms microbes (Kayima et al. 2008; Mbabazi et al., 2010), *Escherichia coli* and *Salmonella* spp. (Fuhrimann et al. 2015) in the Nakivubo Water Channel, highlighting human activities with implications for public health. However, studies on microbial composition and diversity along the Nakibuvo Channel has not been performed. The aim of this study was to gain a better understanding of the microbial diversity changes and response to anthropogenic pollution in the Nakibuvo Water Channel. Consequently, we assessed samples collected from industrial and urban polluted sites by analyzing microbial diversity based on 16S rRNA gene clone-library.

## Materials and Methods

### Sample collection

Water samples were collected from Kibira point [0°18’35.6”N 32°34’31.9”E], Kitante Channel [0°18’57.1”N 32°35’42.5”E] and Nakivubo wetland [inlet 0°18’52.9”N 32°36’36.2”E, main; 0°18’50.2”N 32°36’41.5”E] here referred to as Bugolobi point) along Nakivubo Water Channel, Kampala, Uganda. The channel is a manmade stream that drains Kampala city and its suburbs and empties into Lake Victoria at the Inner Murchison Bay. Kibira and Bugolobi sampling points receive pollutants mainly from low and high impact industrial wastes, respectively; while Kitante channel waste is mainly from urban pollution and human activities.

Genomic DNA was extracted using the MoBio soil extraction kit (MO BIO Laboratories, Carlsbad, USA) following the manufacturer’s recommendations. The extracted DNA was amplified by PCR using 16S rDNA specific primers, visualized by agarose gel electrophoresis and the yield estimated by using ethidium bromide stained agarose gels with lambda DNA as a calibration standard. The 16S rDNA gene amplification products were blunt-end cloned into pUC-19 and used to transform *E. coli* ER2688 cells. Single clones were selected, named based on sampling sites and re-cultured individually. The plasmids were recovered from all the clones and the inserts of approximately 1500 bp sequenced by Sanger method using the universal M13R and M13F primers.

### Data analysis

Clone sequences were edited using CodonCode Aligner v 3.5 (Condon code Corporation, Dedham, USA). All sequences were aligned using ClustalW (align.genome.jp; Larkin et al. 2007). The genetic diversity per locality, as estimated by the number of polymorphic sites, nucleotide diversity (Kimura 2-parameter model), number of haplotypes, and the genetic similarity (i.e. genetic distances by Kimura 2-parameter model) among sampling sites were calculated using DnaSp v5.10 (Librado and Rozas 2009). Operational taxonomic units (OTU) classified at 97% sequence similarity were clustered and the microbial species diversity per sampling site was calculated with Shannon-Weaver diversity index using vegan package (Oksanen et al. 2012) for R. The OTU sequences were submitted to the non-redundant NBCI nucleotide database using Blastn queries (http://blast.ncbi.nlm.nih.gov/) for identification purposes. An e-value cut off (e-5) and >90% sequence similarity were used as characterization thresholds.

Maximum likelihood (ML) trees were constructed per sampling site using all clone sequences in PhyML 3.1 (Guidon and Gascuel 2003) using the default settings under GTR+G model as obtained by jModeltest 3.0 (Posada 2008). The trees were bootstrapped without root using 10000 replications.

## Results and Discussion

The microbial community of river channels and wetlands has served as a monitoring element of aquatic ecosystems as they display particular responses to environmental pollutants (Chakraborty and Bhadury 2015). Here, we present an insight into the microbial diversity changes at industrial and urban polluted sites along Nakivubo Water Channel that drains into Lake Victoria, Uganda. A total of 463 clones of 16S rRNA sequences were analyzed. Clone sequences had an average of 397 bp, 394 bp, and 399 bp for Bugolobi, Kitante and Kibira sampling sites, respectively, representing the 1500 bp of 16S rRNA region. The clones were classified into 35 Operational taxonomic units (OTU). The Blastn identification analysis of our OTU sequences and NBCI database gave a match of 32 bacteria species with > 90% sequence similarity (Table 1). An average of 89% of the OTU within sampling sites matched with the database. A total of 9, 11 and 21 bacteria species were found in Bugolobi, Kitante and Kibira sampling sites, respectively.

**Table 1.**
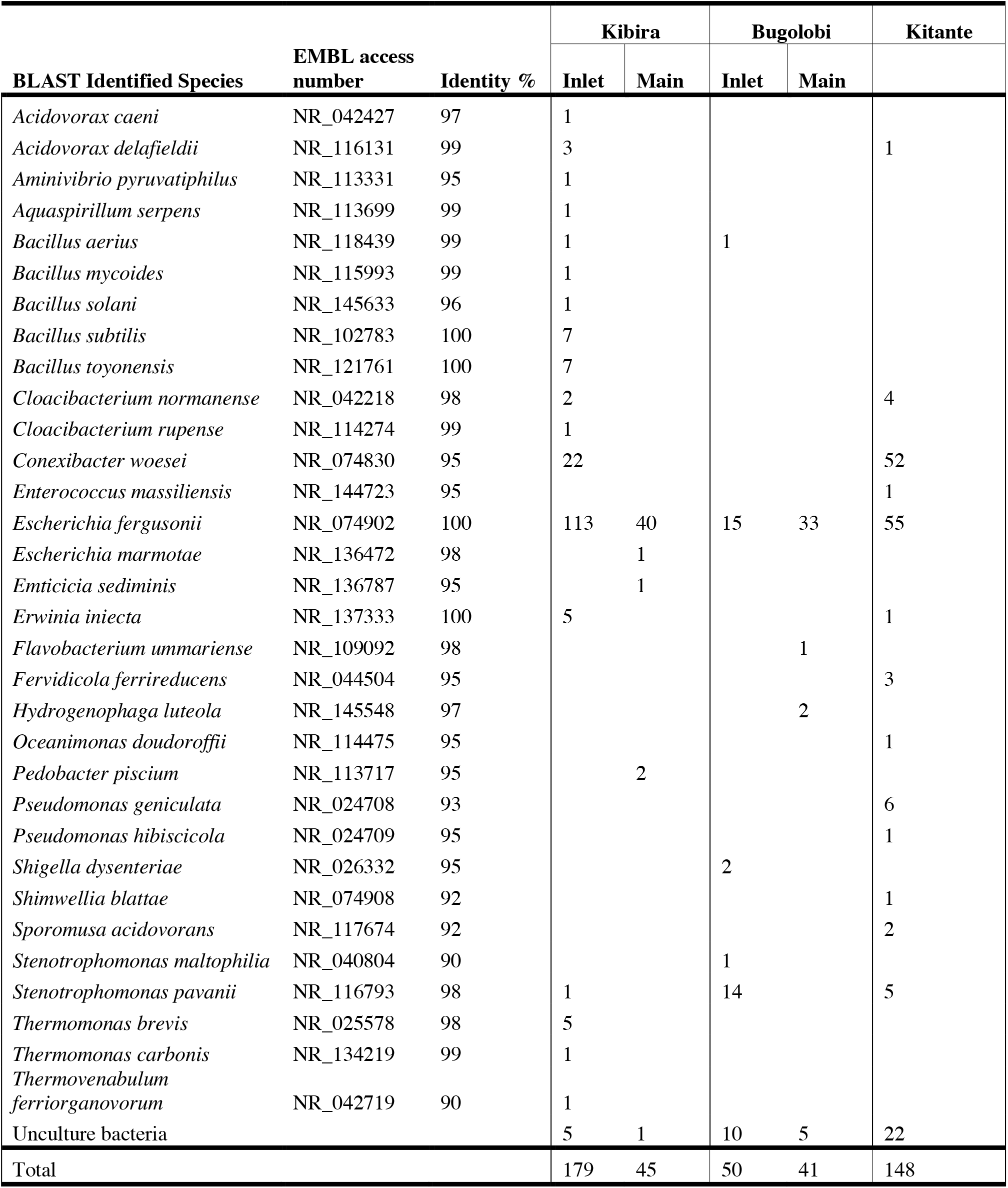
**Comparison of the identity between the operational taxonomic unit (OTU) sequences** from the Nakivubo Water Channel and matched 16S rRNA sequences in the NCBI database (>90% of sequences similarity). The number of matched OTU per sampling sites per locality is shown.

Differences in microbial diversity and community structure were observed at different sampling points (Fig. 1). Shannon diversity index highlighted an increase of microbial diversity (Shannon-Weaver = 0.89), which suggests that industrial pollution influences the adaptation and community distribution of the microbial community, as previously observed (Yang et al. 2020; Aguinaga et al. 2018; Jałowiecki et al. 2016; Ma et al. 2015). The industrial polluted sampling sites (Bugolobi and Kibira) showed high genetic similarity (Kimura 2-parameter model) among microbial profiles than with urban polluted sampling site (Kitante). Among the 28 bacteria species found at the industrial polluted sites, were three unique bacteria, *Bacillus* sp., *Pseudomonas* sp., *Thermomonas* sp., which have been reported to transform contaminants such as heavy metals and hydrocarbons (such as oils) by their metabolic pathways (Bhakta et al. 2014; Kanmani et al. 2012), and are therefore linked to bioremediation (Singh et al. 2014). These microbial communities obtained from the industrial polluted sites of Nakivubo Water Channel could be involved in the biotransformation of pollutants, and therefore could be considered valuable biological indicators of industrial pollution and used for monitoring sewerage and industrial waste treatment processes, and as possible agents of bioremediation in the water channel.

**Figure 1.**
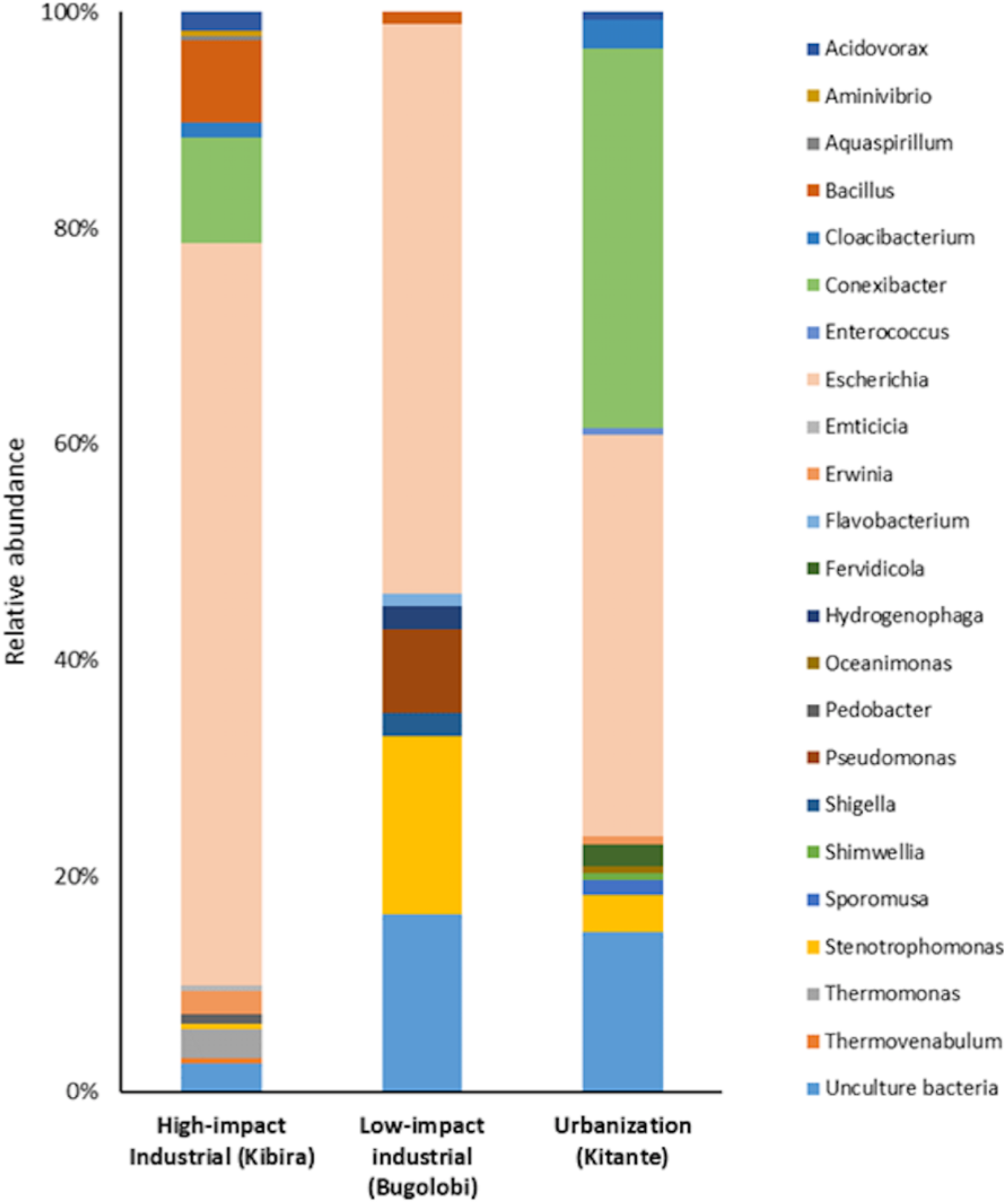
Relative abundance of Nakivubo Water Channel bacterial taxa at the genus level. Each color-bar represents the percentage of the taxa in the total number of clones from each sampling site.

The urban polluted site (Kitante sample site) displayed a low proportion of pathogenic bacteria such as *Enterococcus* sp., *Pseudomonas* sp., and *Stenotrophomonas* sp. The presence of potentially pathogenic bacteria might have adverse public health implications when people come into contact with the water (Egli et al. 2002). These three bacterial groups are associated with urinary tract infections, diarrhea, and pneumonia and other medical conditions as previous reported from other urban polluted sites (Cabral 2010; Egli et al. 2002). Therefore, our findings suggest that control interventions involving channel sanitation should be undertaken in order to mitigate health risks to the public.

*Escherichia* sp. was the most common and abundant microorganism for both industrial and urban polluted sites at Nakivubo Water Channel. High concentrations of *E. coli* was also previously observed in the Channel (Kayima et al. 2008; Fuhrimann et al. 2015). *E. coli* sp. strains can cause gastroenteritis, diarrhea, and vomiting in humans, among other infections, by contact or drinking contaminated water (Cabral 2010). Therefore, human access to channel water should be restricted and continuous monitoring of the channel implemented.

Lastly, the aligned clone sequences contained an average of 84 polymorphic sites, 61 parsimony-informative sites, 23 singletons, and 0.68 of nucleotide diversity per locality (Table 2). The phylogenetic reconstruction of maximum likelihood trees (Bugolobi, likelihood: logL = −3733.3; Kitante, likelihood: logL = −3462.0; and Kibira, likelihood: logL = −6085.8) showed different clustering results in congruency with the OTU cluster analysis and the matched results with the NCBI database (Fig. 2). Bugolobi showed a branch separation of *Escherichia* species sequences based on main and inlet sampling points. These results suggest that two possible different *Escherichia* strains co-habit the channel as previously reported (e.g. Chekabab et al. 2013) and should be further monitored. Both Bugolobi and Kibira sampling sites showed that the inlet sampling sites were more genetically diverse than the main sampling site, thereby suggesting the existence of distinct sources of pollution to the site.

**Figure 2.**
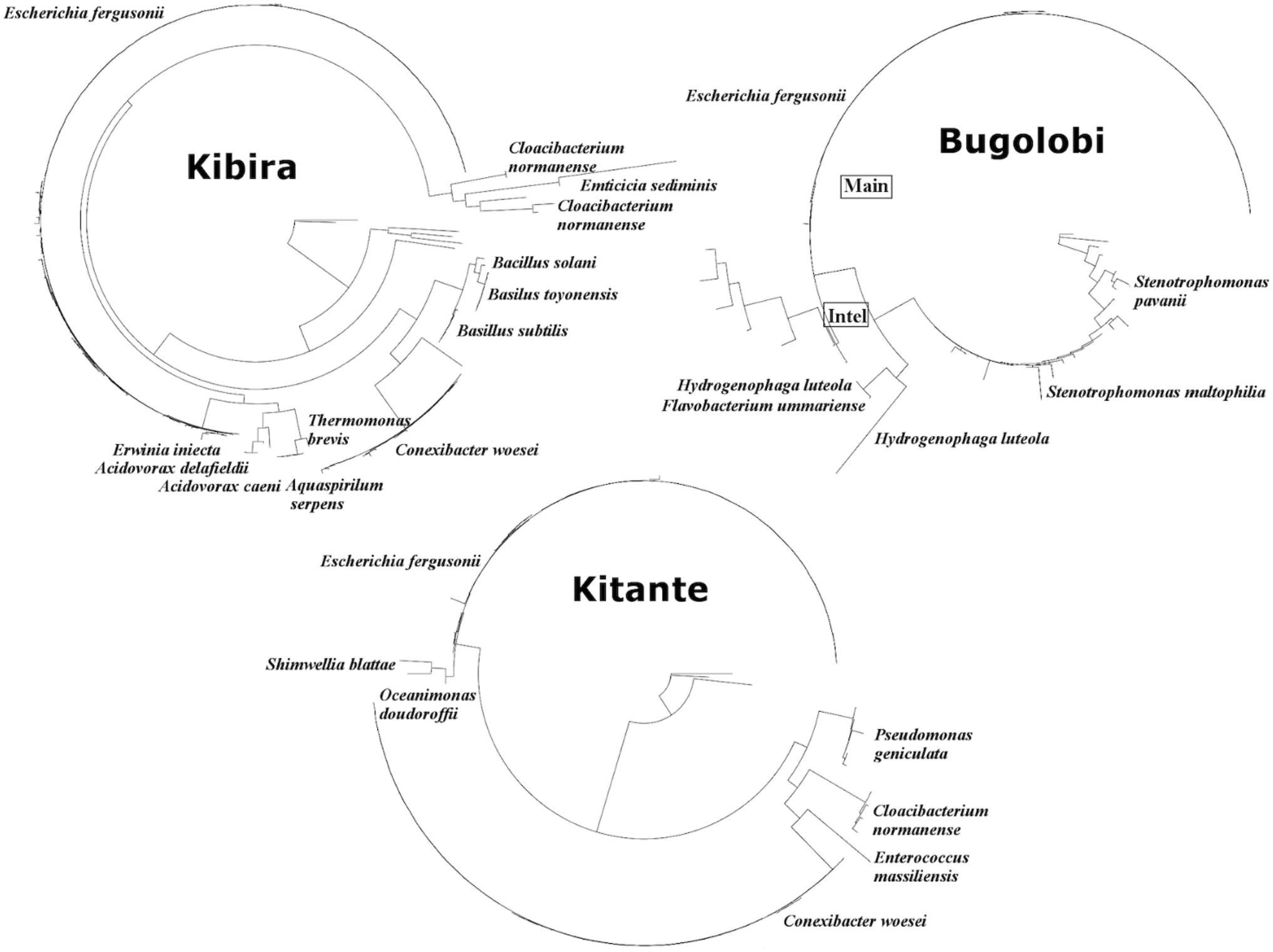
Maximum likelihood phylogenetic trees based on the 16S rRNA gene sequences from three sampling sites: Kibira, Bugolobi, and Kitante along the Nakivubo Water Channel, Uganda. Sequences identified in the NCBI database were added to the phylogenetic trees. Branches that displayed values below 50 were collapsed.

**Table 2.**
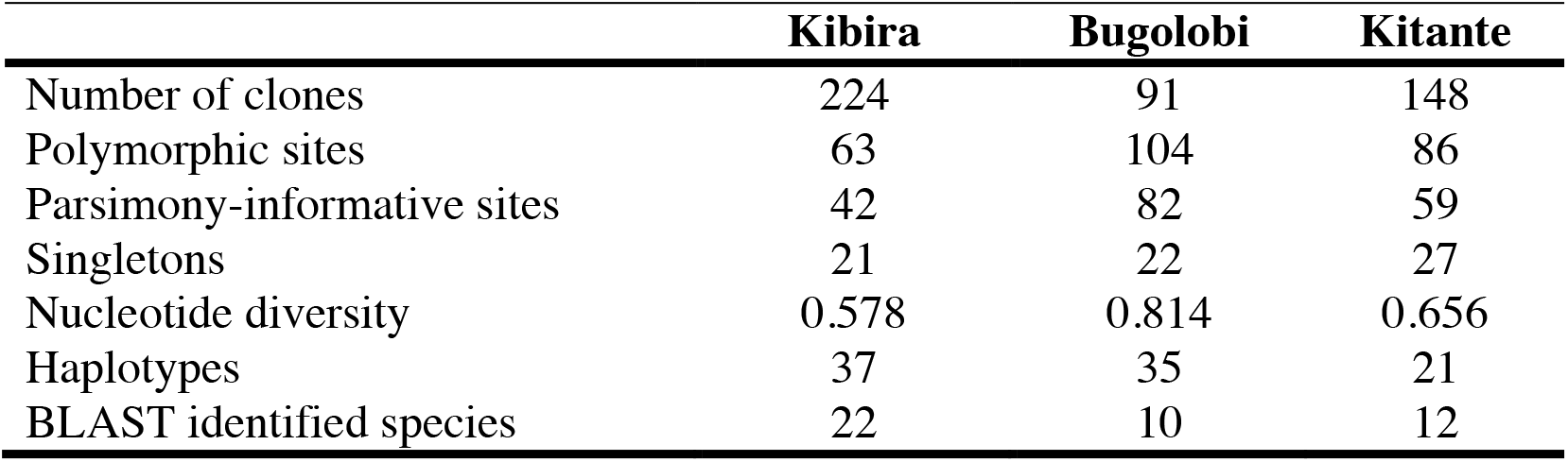
Summary of the genetic diversity per locality.

|  | **Kibira** | **Bugolobi** | **Kitante** |
| --- | --- | --- | --- |
| Number of clones | 224 | 91 | 148 |
| Polymorphic sites | 63 | 104 | 86 |
| Parsimony-informative sites | 42 | 82 | 59 |
| Singletons | 21 | 22 | 27 |
| Nucleotide diversity | 0.578 | 0.814 | 0.656 |
| Haplotypes | 37 | 35 | 21 |
| BLAST identified species | 22 | 10 | 12 |

## Conclusions

Microbial communities provide significant resolution for in-depth monitoring of ecosystems health and extremely useful analytical systems for water pollution. We showed that potential bacterial adaptation could be associated with industrial pollution and could play a role for future bioremediation of the channel while pathogenic bacteria were associated with urban pollution. Both polluted sites showed a high concentration of *E. coli* that may pose risks to human health. The observed cumulative response of microbial communities to pollutant exposure can be leveraged to improve monitoring and pollution assessment. In addition, increased targeted wastewater treatment before dumping into the Channel, as previously suggested (Kayima et al. 2008; Fuhrimann et al. 2015; Matagi 2002; Mbabazi et al. 2010; Emerton 1998) could improve water quality and restore natural microbial communities. The diversity of natural microbial consortia could also be harnessed as a rich source for novel industrial enzymes as well as medicinal and bioactive compounds

## Acknowledgments

We thank the research teams at Med Biotech Laboratories for technical support and Professor Francis Mulaa (University of Nairobi, Kenya) for constructive suggestions and discussions.

